# The diurnal transcriptional landscape of the microalga *Tetradesmus obliquus*

**DOI:** 10.1101/425934

**Authors:** Benoit M. Carreres, G. Mitsue León-Saiki, Peter J. Schaap, Ilse M. Remmers, Douwe van der Veen, Vitor A.P. Martins dos Santos, René H. Wijffels, Dirk E. Martens, Maria Suarez-Diez

## Abstract

*Tetradesmus obliquus* is a promising oleaginous microalga. We functionally annotated its genome and characterized the transcriptional landscape of *T. obliquus* adapted to 16:8h light dark (LD) cycles in turbidostat culture conditions at very high temporal resolution (1h intervals). Revealing a cycle of cellular events, six distinct expression profiles were obtained, each with transcriptional phenotypes correlating with measurements of biochemical composition.

The impact of starch deficiency was studied using the starchless mutant *slm1*. Significant changes in the transcriptional landscape were observed. Starch deficiency resulted in incapacity to supply energy during dark period, resulting in early or late time shift for energy demanding processes. Our study provides new perspectives on the role of starch and the adaptation to LD cycles of oleaginous microalgae.

## 1 Introduction

Microalgae are a promising source of compounds of interest (lipids, proteins, and pigments) for the production of food, feed, chemicals, and fuels [1–3]. Large scale microalgal production will be primarily done outdoors under natural diurnal light/dark (LD) cycles [4, 5]. Diurnal cycles are ubiquitous and photosynthetic organisms synchronize their metabolic activities to anticipate light changes in the environment and schedule specific tasks during the night or day [6–9]. Synchronization in photosynthetic organisms involves the regulation of photosynthesis to maximize carbon fixation and use of light during the day and to schedule light sensitive processes (such as DNA synthesis and cell division) at night [6, 10–12].

*Tetradesmus obliquus* is a microalga recognized as an industrially relevant strain for food and fuel production [13, 14]. *T. obliquus* can reach a maximum triacylglycerides (TAG) content of 0.45 g⋅g_DW_^−1^ and a maximum TAG yield on light of 0.14_g⋅_mol_ph_^−1^ under batch nitrogen starvation and continuous light conditions [15]. For further improvement of TAG yield on light, de Jaeger et al. developed the starchless mutant *slm1* [16], which cannot synthesize starch due to a missense mutation in the small subunit of ADP-glucose pyrophosphorylase, the committed step of starch biosynthesis [17]. Under culture conditions of continuous light and batch nitrogen starvation, *slm1* showed a higher maximum TAG yield on light (0.217 g⋅mol_ph_^−1^) and maximum TAG content (0.57 g⋅g_DW_^−1^) compared to the wild-type (WT) without a decrease in photosynthetic efficiency [15]. Comparable results were obtained under light-dark cycles and batch nitrogen starvation [14].

LD cycles give an advantage over continuous light, leading to higher energy conversion efficiency and higher yield of biomass on light. The physiological behavior of *T. obliquus* WT and of the starchless mutant (*slm1)* was also studied under 16:8h LD cycles in turbidostat controlled systems [8, 10]. Under this light regime and nitrogen replete conditions, *T. obliquus* showed synchronization and diurnal patterns of its metabolism, which amongst others, suggested that starch was used as a temporary energy storage. This is, starch was accumulated during the light period and was used during the dark period. Cell division was also synchronized and occurred mainly during the night. The starchless mutant *slm1* showed a lower energy conversion efficiency (11-24% lower) and biomass yield on light (13-39% lower) compared to the WT under different photoperiods [10]. Furthermore, for the *slm1* mutant cell division still occurred mainly during the night, but at a slower rate and no diurnal oscillations on any of the other measured compounds were found [10]. Unlike for the WT, diurnal LD cycles did not provide an advantage for the *slm1* mutant compared to continuous light. On light, biomass yield as well as the energy conversion efficiency were similar [8]. Starch may therefore play an important role in harvesting additional light energy during LD cycles.

While the biochemical analysis allowed us to draw some conclusions, we still lack information on how cellular processes are regulated during the diurnal cycle and how the synthesis and use of starch is connected to these processes. Furthermore, while the inability to make starch had an impact on energy conversion efficiency, it is not known how this will affect the timing and regulation of the different cellular processes. LD cycles and the subsequent synchronization of the microalgal population makes it of paramount importance to unravel the timing of cellular and subcellular events. Understanding the timing at which metabolic changes take place is essential to understand the phenotype exhibited by the WT and *slm1* strains and to optimally design experiments characterizing mutant strains.

Time resolved transcriptome analysis can give useful insights into the timing and regulation of the different physiological stages and on the succession of processes in the cell. Thus it allows the association of the cellular processes with biochemical properties. Algal diurnal transcriptional regulation is not well known. To our knowledge, only a few reports on diurnal oscillations under light/dark cycles in microalgae have been published [18–23] The first one was on the eukaryotic red alga *Cyanidioschyzon merolae* [24] grown under 12:12h LD cycles, where they studied the transcriptional changes in intervals of 2 h. Furthermore, the diurnal transcriptional changes were studied in the diatom *Phaeodactylum tricornutum* [20] under 16:8h LD cycles. The authors studied changes in biochemical composition (carbohydrates and lipids) in 8 different unequally distributed time points during a period of 26.5 hours (5 points during the light period and 3 during the dark period). Finally, transcriptional analysis on *Nannochloropsis oceanica* [21] and *Chlamydomonas reinhardtii* [22] under a 12:12h LD cycle was published. For *N. oceanica* the authors studied growth, changes in biomass composition (lipids and glucose) and in gene expression in intervals of 3 h., while for *C. reinhardtii* a more detailed study was done with intervals of 1h during most of the cycle and every 30 min for some time points, ending up with a total of 28 points distributed over the 24h cycle. Our study, however, is the first one to look into the transcriptional changes in a diurnal cycle of an oleaginous green algae with a high sampling frequency (intervals of 1 h). Additionally, this is the first study reporting on the transcriptome changes over a diurnal cycle for a starchless mutant, which could give insights into the role of starch in microalgae.

To fully characterize the changes induced by LD cycles and the role of starch metabolism, we analyzed and compared the transcriptional landscape of *T. obliquus* WT and *slm1* under diurnal 16:8h LD cycles. For this, we cultivated both strains in continuous turbidostat controlled photobioreactors. Samples were taken after the oscillating steady-state was synchronized to the LD cycle. To obtain a high resolution for WT, sampling was done in intervals of 1h, while for *slm1* sampling was done at 3h intervals.

## 2 Materials and methods

### 2.1 Experimental setup and sampling

Wild-type (WT) *Tetradesmus obliquus* UTEX 393 (formerly known as *Acutodesmus obliquus and Scenedesmus obliquus* [25]) was obtained from the Culture Collection of Algae, University of Texas. The starchless mutant of *T. obliquus* (*slm1*) was generated as described by de Jaeger et al. [16]. *T. obliquus* was continuously cultivated in a sterile flat panel airlift-loop reactor with a 1.7 L working volume (Labfors 5 Lux, Infors HT, Switzerland). Culture conditions and reactor set-up (27.5 °C, pH 7.0 and gas flow rate of 1 L⋅min^−1^ air enriched with 2% CO2) were controlled as described by León-Saiki et al. [8]. Light was provided in a 16:8h light/dark (LD) block at an incident photon flux density of 500µmol⋅m^−2^⋅s^−1^ (warm white spectrum 450-620 nm). Cultivations were turbidostat controlled, where fresh medium was fed to the cultures when the light intensity at the rear of the reactor dropped below the setpoint (10µmol⋅m^−2^⋅s^−1^, OD_750_). Feeding of medium was stopped during the dark period. After steady state was reached, 8mL samples were taken for RNA extraction (approximately 10 mg_DW_). Cells were immediately collected by centrifugation (4255 xg, 0°C for 5 min), supernatant was discarded and pellets were frozen in liquid nitrogen and stored at −80°C until further extraction. Samples for RNA extraction were taken in intervals of 1h for WT or every 3h for *slm1*. Due to restrictions on working hours of the laboratory, the samples were collected in two successive time settings to allow sampling the dark period during the day. After collecting samples of the first half of the cycle, light settings were shifted and the culture was then allowed to reach oscillating steady-state before collecting samples for the second half of the cycle. The first and the last samples of each time settings are overlapping samples for control. Therefore, four RNA samples are present at these time points. Overall 72 samples were taken for RNA extraction.

### 2.2 RNA isolation and quality control

RNA extraction was performed using the Maxwell® 16 LEV simplyRNA Tissue kit (Promega). Frozen algae pellet (≈200 μL) were submerged in 400 μL of homogenizing buffer supplemented with 8 μL 1-thioglycerolL) were submerged in 400 μL) were submerged in 400 μL of homogenizing buffer supplemented with 8 μL 1-thioglycerolL of homogenizing buffer supplemented with 8 μL) were submerged in 400 μL of homogenizing buffer supplemented with 8 μL 1-thioglycerolL 1-thioglycerol in a 2 mL Lysing matrix C tube (MP), prefilled with a mix of glass beads. Samples were disrupted using a FastPrep-24 instrument (MP). After disruption, all liquid was transferred to a LEV RNA Cartridge. 300 μL) were submerged in 400 μL of homogenizing buffer supplemented with 8 μL 1-thioglycerolL lysis buffer (from the kit) were added and the rest of the extraction was performed using a Maxwell MDx AS3000 machine (Promega) following manufacturer’s instructions. RNA integrity and quantity were assessed with an Experion system (Bio-Rad), and only high quality samples (RIN value ≥ 7) were selected. Total RNA was sent for whole transcriptome sequencing to Novogene Bioinformatics Technology Co. Ltd (HongKong, China).

The corresponding data have been submitted to EBI ArrayExpress and can be found under the accession number E-MTAB-7009.

### 2.3 Genome structural annotation

Using the available genome sequence of *T. obliquus* [26], we performed an RNA-Seq-based genome annotation using BRAKER1 [27]. RNA reads from 38 samples of both WT and *slm1*, supplemented with an additional sample from each strain under nitrogen limited condition were given as additional BRAKER1 input. The annotation information was processed using our semantic framework pipeline [28] and the information was stored according to our integrated ontology [29] which respects the FAIR data principles [30]. The annotation framework and the ontology were extended with the necessary tools [27, 31, 32] and ontology terms for the purpose of this analysis.

The genome feature annotation has been used to updated the original genome ENA project with accession number PRJEB15865.

### 2.4 Genome functional annotation and pathway mapping

Proteins were annotated to GO terms using InterProScan5 and Argot2, with default parameters [33, 34]. Proteins were annotated to enzyme commission (EC) number using EnzDP [35]. Results with a complete EC number and a likelihood-score of at least 0.2 were used for subsequent analysis. To choose a threshold for the likelihood-score that results in a good trade-off between true positive and false positives, we visually compared the completeness of KEGG metabolic maps while avoiding dispersion of each reaction into different expression clusters. The functional annotation can be found in supplementary file S1 that includes EC numbers and GO terms.

### 2.5 RNA-seq normalization and expression calculation

Using all the transcripts found from the genome annotation, we aligned the reads from each sample and calculated the FPKM values using Cufflinks [32]. As a pre-filter to remove unexpressed genes and false positives from gene annotation, we selected the transcripts with a coverage of at least 1 and a FPKM of at least 0.1 in at least 10 of the 113 analyzed samples. These samples include the 72 of this study, and the rest were taken under LD cycles and nitrogen limited conditions (not studied in this manuscript). Additionally, we selected the transcripts having an expression of at least 10 samples with a value higher or equal to the 0.15 quantile. Following the advice of maSigPro [36], the FPKM of all samples were normalized using the scaling normalization method TMM [37] using the R functions from edgeR package “calcNormFactors” [38].

### 2.6 Gene clustering

To identify genes with significant expression profile changes over time, maSigPro was used [36, 39], with the modified parameters: regression model was set to a maximum of 23 degrees, the parameter “counts” was set to true, “nvar.correction” was also set to true, the “step method” set as backward, and R-squared was set to 0.7. The R-squared was chosen based on the relevance and the balance in the number of enriched pathways identified in each cluster (see below). To better estimate the number of clusters to group the gene expression, we evaluated combinations of significance level (Q: 0.005 to 0.05) and R-squared (0.85 to 0.5). We decided to keep the standard-strict values (Q=0.05, R-squared=0.7), which would also give a good balance in pathway enrichment and number of genes in each group.

Hierarchical clustering with agglomerative linkage was performed using the R stats package (hclust function). Number of clusters from 3 to 25 were evaluated using the R function “cluster.stats” from package “fpc” [40]. For each set of clusters, the following indexes were computed: average silhouette widths, normalized gamma, two dunn indexes, average within and average between ratio, and Calinski and Harabasz index. Additionally, each set of clusters were compared one on one with the Rand index.

### 2.7 Enrichment analyses

Enrichment analyses were performed using the hypergeometric function to model the probability density using the “phyper” function from the R package stats [41]. Two types of analysis were performed: pathway and GO term enrichment. Pathway enrichment required associating annotated Enzyme Commission (EC) numbers to metabolic maps. We used the online available resource from KEGG pathway maps [42, 43]. The KEGG pathways fitting the following requirements were kept for further analysis: 60% coverage if 3 to 6 EC numbers annotated, and 50% coverage if 6 to 10 EC numbers annotated, and 25% of coverage if more than 10 EC numbers were annotated. For the hypergeometric test we considered the universe size, *N*, to be the total number of EC numbers in all pathways in the genome, *m* is the number of successes in this universe and is defined as the number of EC numbers in the corresponding pathway in the genome, *k* and *x* are the sample size and the number of successes in the sample (or considered gene subset) respectively. Enrichments with a p-value lower than 0.05 were considered significant. Similarly, for the GO enrichment, *N* is the total number of genes annotated to any GO terms in the genome, *m* is defined by the number of genes annotated to the considered GO term in the genome, and *k* and *x* refer to the considered subset of genes. Multiple test correction for the GO enrichment was performed using the Benjamini–Hochberg procedure. Enrichments with FDR<0.05 were considered significant. To handle the GO information such as ontology, ancestor, and offspring, we used “GO.db” database from Bioconductor [44]. Additionally, to reduce the number of GO terms and conserve the most specific, only terms not having any offspring in the selection were retained. We additionally provide the pathway coverage, and the *x* and *m* values to better understand the reason of each significant enrichment.

## 3 Results

### 3.1 Genome annotation

The genome of *T. obliquus* UTEX 393 is about 100 million base pairs in size [26]. BRAKER1 annotation revealed 19795 genes, which transcribed 21493 coding sequences and translates into 19723 protein sequences.

To analyze the changes in transcription over a diurnal cycle of *T. obliquus* WT and *slm1* strains, we sequenced RNA from biological duplicate runs. WT was sampled every hour in duplicate resulting in 52 samples. *slm1* was sampled in intervals of three hours resulting in 20 samples. For both strains, time points 0h and 13h were overlapping samples and therefore sequenced in quadruplicate (see material and methods for details). Not all the transcripts were found to be expressed and filtering was performed to identify genes with very low expression that were not further considered. From the 19723 proteins, 16810 proteins remained after filtering for subsequent analysis. GO term annotation yielded 13687 proteins annotated to 2394 unique GO terms. EC annotation yielded 3559 proteins annotated to 1315 EC numbers. The functional annotation can be found in supplementary file S1.

### 3.2 Light-dark 16:8h cycle induces systemic transcriptional changes following a circular pattern

4686 genes were found to have a significant change of expression during the diurnal cycle. A principal component (PC) analysis (PCA) for these selected genes is shown in Figure 1A. The PC plot provides a global overview of changes in gene expression over time. The time point samples form an ordered and circular pattern over the two first PCs. These two first PCs together explain 87% of the variation in expression. To demonstrate that this is not a consequence of gene selection, we rendered another PCA considering the whole set of expressed genes (supplementary file S2). Within this set, the overall pattern is noisier, but the two first components still explain 78% of the variation in expression. The main changes in gene expression occur along PC1 from 1h to about 5h and back again from 8h to about 13h. The second important change occurs along PC2 from 6h to 9h and back again from about 22h to 0 h. During the period from 14h to 22h there is very little change in gene expression. Overall, genes associated to light harvesting complex (LHC) were found to be the main contributors of PC1. The two highest contributing genes to PC2 are glyceraldehyde-3-phosphate dehydrogenase (GA3PDH) and glucose-6-phosphate isomerase (GPI). LHC genes also contributed strongly to PC2.

**Figure 1:**
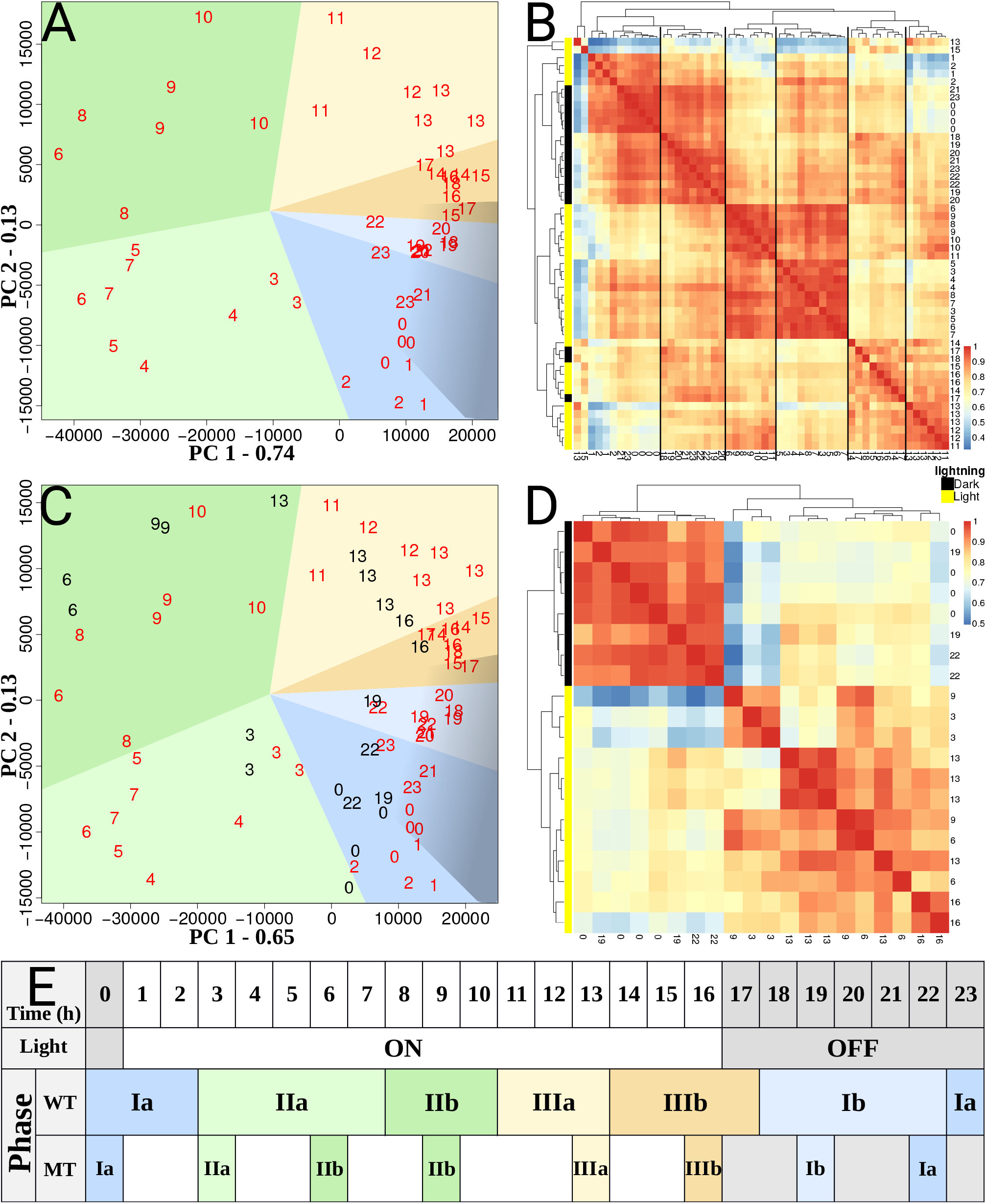
Principal Component Analysis (PCA) (A,C) of gene expression data for Tetradesmus obliquus wild-type (WT) (A) and both WT and starchless mutant (slm1) (C) during 16:8h light-dark cycle. Numbers represent the time points of the samples, with red for the WT samples and black for the slm1 samples. Background colors in both PCA plots refer to the colors given to the six time sub-phases described in (E), and the dark shade refers to the dark period. (B, D) Heatmaps showing similarity of gene expression data among samples for WT (B) and slm1 (D) strains, samples are identified by their time points. Dark vertical lines across the heatmap in (B) indicate different time time phases. (E) Overview of the time phases.

The heatmap representation in Figure 1B offers a complementary view of these multi-dimensional data. It allows to evaluate differences and similarities in expression between the time points. Two samples corresponding to time points 13h and 15h are separated in the dendrogram from the neighboring samples and from the biological duplicate samples. However, in the heatmap it can be seen that these samples still show relatively high correlation with their corresponding biological duplicates and samples from the surrounding hours. Three main groups of time points can be identified in the dendrograms in Figure 1B, which we will refer to as time phases I, II, and III. Additionally, each phase can be subdivided in two sub-phases, a and b, which represent subtler expression changes. The occurrence of the phases over the 16:8h LD cycle is shown in the Figure 1E for both strains. The phases were named according to their order of appearance from 0h to 23h. These time phases are also apparent and can be associated to changes along the two first PC. Similarly, time phases were also associated to the *slm1* strain based on the PCA plots. For *slm1*, we noticed a change in phase timing that represents a significant change of expression occurring one to two hours before each dark-light shift.

### 3.3 Gene expression clusters and their time profiles

#### 3.3.1 Cluster profiles

We clustered the 4686 genes that showed significant changes over time in different numbers of clusters ranging from 3 to 25. Next we evaluated the cluster separation using seven well established indexes to assess similarities within clusters and differences between clusters. These results are depicted in the supplementary file S3. Based on these results, we considered separating the genes in six clusters to be optimal.

The resulting six cluster profiles were summarized using for each time point the median value of gene expression of each gene present in the cluster and are depicted in Figure 2. Cluster 1 to 6 contain respectively 829, 364, 1020, 929, 952 and 592 genes. All the cluster profiles are different, but there are certain similarities between clusters 1 and 6, and between 3, 4, and 5. Interestingly, the two peaks of expression observed in cluster 3, 4, and 5 are all 5-6 hours apart.

**Figure 2:**
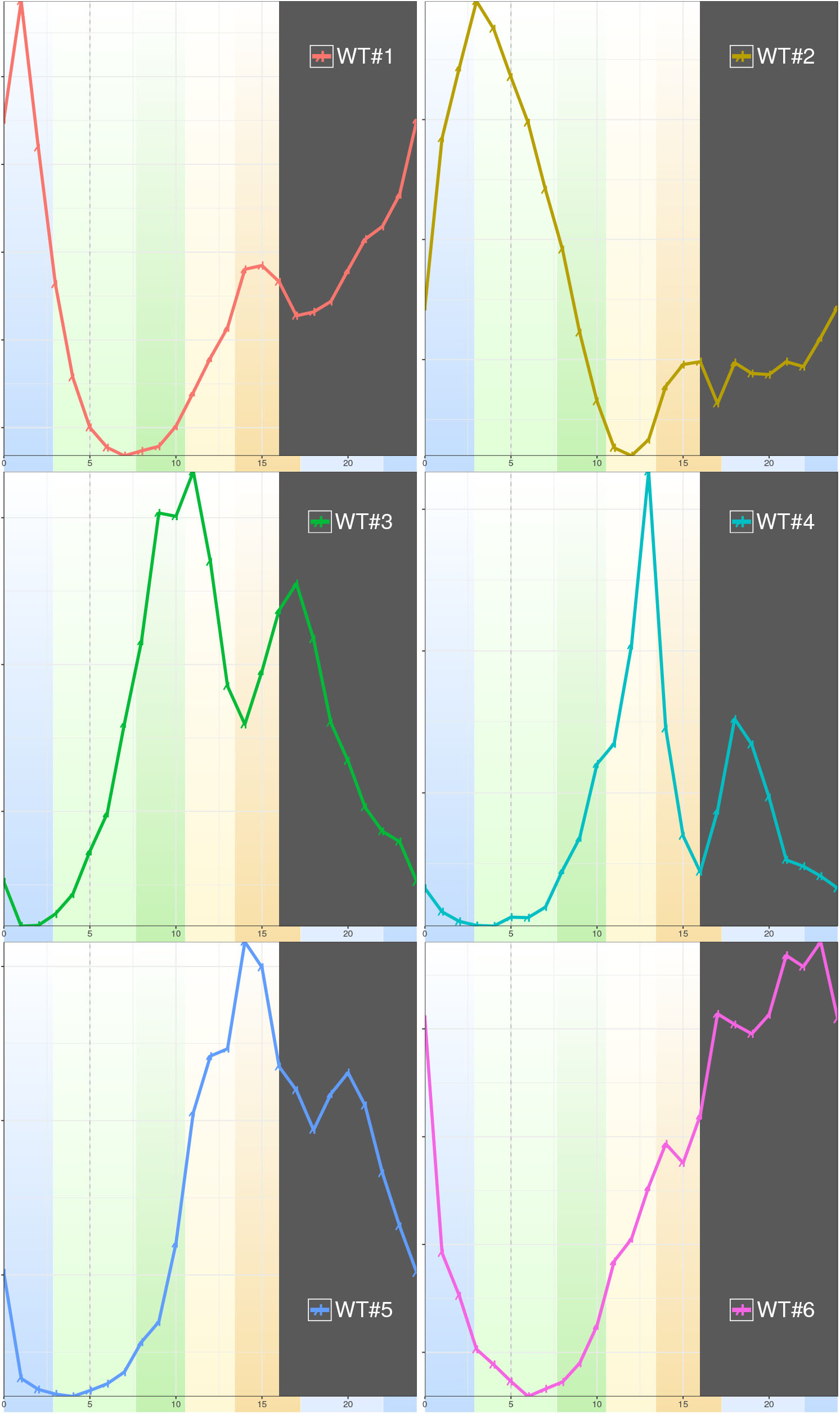
Expression profile of the 6 gene clusters for Tetradesmus obliquus under 16:8h light-dark cycles. The plots show the median profile of gene expression in the indicated clusters. The background colors correspond to the colors given to the time phases in Figure 1. The vertical gray dashed line represents the time point of maximum dilution rate taken from [8]. Dark area represents the dark period.

The temporal profile of each cluster peaks at different moments of the diurnal cycle. Cluster 1 peaks the earliest, 1h after light goes on. Cluster 2 peaks at 3h, cluster 3 peaks around 11h and shows a smaller secondary peak at 17h. Cluster 4, with a similar profile, shows the first peak at 13h and the second at 18h. Cluster 5 has a rather gradual increase of expression and peaks around 14h followed by a smaller peak at 20h. Finally, cluster 6 has very gradual expression changes throughout the day, and shows a peak at 23h.

#### 3.3.2 Cluster functional analyses

To understand the functions of the genes in the clusters, we performed two sets of complementary enrichment analyses. Metabolic pathway enrichment analysis provides direct information on gene metabolic functions. The functional annotation identified genes associated to EC numbers. Mapping these EC numbers to KEGG pathway maps revealed 67 pathways for which enough genes can be associated to the corresponding reactions to warrant further analysis, as detailed in Materials and methods. GO enrichment analyses were performed on the three ontologies: biological process (BP), molecular function (MF) and cellular component (CC). These analyses provide a wider overview not restricted to metabolism. The enrichment results are summarized in Table 1. Each cluster median time profiles and the full set of significant (p-value < 0.05) enrichments are presented in the supplementary file S4.

**Table 1:** Summary of the results of the enrichment analyses. The first column contains the cluster identifier. The second contains the enriched pathways (p-value < 0.0.5). The last three columns are the results of the GO enrichment analyses (FDR<0.05) for each of the GO ontologies: biological process (BP), cellular component (CC) and molecular function (MF). Full set of GO enrichment results are available in the supplementary file S4.

| Cluster | Pathways | BP | CC | MF |
| --- | --- | --- | --- | --- |
| 1 | Ubiquinone and other terpenoid-quinone biosynthesis<br>Purine metabolism<br>Glycine, serine and threonine metabolism<br>Cysteine and methionine metabolism<br>Valine, leucine and isoleucine biosynthesis<br>Lysine biosynthesis<br>Arginine and proline metabolism<br>Riboflavin metabolism<br>Aminoacyl-tRNA biosynthesis | RNA methylation<br>tRNA modification<br>translation<br>nucleobase-containing small molecule metabolic process<br>alpha-amino acid biosynthetic process | ribosome | RNA binding<br>structural constituent of ribosome<br>GTPase activity<br>GTP binding<br>S-adenosylmethionine-dependent methyltransferase activity<br>ligase activity, forming carbon-nitrogen bonds |
| 2 | Glycine, serine and threonine metabolism<br>Phenylalanine, tyrosine and tryptophan biosynthesis<br>Carbon fixation in photosynthetic organisms<br>Porphyrin and chlorophyll metabolism<br>Terpenoid backbone biosynthesis<br>Carotenoid biosynthesis<br>Aminoacyl-tRNA biosynthesis | tRNA aminoacylation for protein translation<br>steroid biosynthetic process<br>coenzyme metabolic process<br>porphyrin-containing compound biosynthetic process<br>photosynthesis<br>pigment biosynthetic process<br>oxidation-reduction process | photosystem | nucleotide binding<br>3-beta-hydroxy-delta5-steroid dehydrogenase activity<br>aminoacyl-tRNA ligase activity<br>iron ion binding<br>calcium ion binding<br>oxidoreductase activity, acting on a sulfur group of donors<br>oxidoreductase activity, acting on paired donors, with incorporation or reduction of molecular oxygen<br>transferase activity, transferring alkyl or aryl (other than methyl) groups<br>carbon-carbon lyase activity<br>hydro-lyase activity<br>heme binding<br>precorrin-2 dehydrogenase activity<br>coenzyme binding<br>2 iron, 2 sulfur cluster binding |
| 3 | Lysine degradation<br>Phenylalanine metabolism<br>Glutathione metabolism<br>N-Glycan biosynthesis<br>Various types of N-glycan biosynthesis<br>Propanoate metabolism | DNA replication<br>DNA repair<br>DNA recombination<br>protein complex assembly<br>vesicle-mediated transport<br>cell cycle process<br>regulation of cellular metabolic process<br>proteasome-mediated ubiquitin-dependent protein catabolic process<br>organelle fission<br>chromosome organization<br>regulation of primary metabolic process<br>carbohydrate derivative biosynthetic process<br>single-organism organelle organization | microtubule<br>cytoskeleton<br>protein complex<br>cytoskeletal part | DNA binding<br>DNA-directed DNA polymerase activity<br>ATP binding<br>microtubule binding<br>transcription factor binding<br>four-way junction helicase activity<br>ATPase activity, coupled |
| 4 | Galactose metabolism<br>Starch and sucrose metabolism<br>Other glycan degradation<br>Amino sugar and nucleotide sugar metabolism<br>Metabolism of xenobiotics by cytochrome P450<br>Drug metabolism - cytochrome P450 | carbohydrate metabolic process |  | hydrolase activity, hydrolyzing O-glycosyl compounds<br>microtubule binding<br>polysaccharide binding |
| 5 | Glycolysis / Gluconeogenesis<br>Glutathione metabolism<br>Starch and sucrose metabolism<br>Sphingolipid metabolism<br>Pyruvate metabolism<br>Glyoxylate and dicarboxylate metabolism<br>Thiamine metabolism<br>Pantothenate and CoA biosynthesis<br>Nitrogen metabolism<br>Metabolism of xenobiotics by cytochrome P450<br>Drug metabolism - cytochrome P450 | carbohydrate metabolic process<br>oxidation-reduction process |  | oxidoreductase activity, acting on CH-OH group of donors |
| 6 | Fatty acid degradation<br>Pyrimidine metabolism<br>Butanoate metabolism<br>Folate biosynthesis |  |  |  |

Cluster 1 shows enrichment in pathways related to amino acids (AA), and riboflavin metabolism. GO terms enrichment is in agreement and contains terms such as tRNA modification, aminoacyl-tRNA biosynthesis, ribosome, alpha-amino acid biosynthetic process and translation. All these pathways and terms indicate protein synthesis and the associated strong demand for AA.

Cluster 2 mainly shows enrichment in the processes related to pigments synthesis, which include carotenoids, chlorophylls, and their precursor (Phytyl-diphosphate) from the terpenoids backbone synthesis. Those are needed for building the photosystems and starting the carbon fixation. Cluster 2, also shows enrichment in genes related to transcription, amino acid (AA) and protein synthesis, but to a lower extent than cluster 1. The carbon fixation pathway is found enriched in cluster 2, which goes in line with enrichment in the pigments and in starch synthesis. The GO terms in biological process are in agreement with the pathway enrichments. The GO terms in molecular function provide extra information on the nature of chemical reactions performed by these genes. We also found that this cluster contains genes associated to the LHC that were found to strongly contribute to PC1 and PC2. Interestingly, while all the other amino-acid synthesis pathways were found enriched in cluster 1, the pathway of “Phenylalanine, tyrosine and tryptophan biosynthesis” is found enriched in cluster 2. This result indicates that these aromatic amino-acids are probably synthesized later than the other, more simple, amino-acids.

In cluster 3, the GO enrichment provides valuable information that cannot be covered by the pathway enrichment. The GO term enrichments display clear terms such as “DNA replication”, “organelle fission”, “chromosome organization”, “microtubule”, “cytoskeletonDNA-directed”, “DNA polymerase activity”. This cluster clearly groups all processes related to the full cell cycle. The pathway enrichment reveal processes related to AA pathways: “Lysine degradation” and “Phenylalanine metabolisms”. Additionally, the enrichment in “N-Glycan biosynthesis” and “Various types of N-glycan biosynthesis” together with the enriched GO terms “protein complex assembly” and “vesicle-mediated transport” indicates important protein maturation processes leading to complex proteins and some transport of proteins to membranes.

Cluster 4 is mostly enriched in pathways associated to different kind of carbohydrates metabolism. The first pathways enriched are “Galactose metabolism“, “Starch and sucrose metabolism”, “Other glycan degradation”, and “Amino sugar and nucleotide sugar metabolism”. With the GO terms displaying “polysaccharide binding”, “hydrolase activity”, and “hydrolyzing O-glycosyl”, we observed the same trend towards starch degradation. The GO term for “microtubule binding” is the only one of this kind in this cluster, but it comes right after cluster 3 where a lot of cytoskeletal and microtubule terms were found enriched. Finally, there is enrichment in Cytochrome P450 which hints on repair mechanisms potentially related to photo-damage.

Cluster 5 shows enrichment in a very diverse set of metabolic pathways, covering glycolysis, pyruvate metabolism, glutathione metabolism, co-enzyme-A, starch and other carbohydrates polymers. More importantly, the cluster is also enriched in genes related to nitrogen metabolism possibly linked to nitrogen assimilation.

Cluster 6 shows enrichment in fatty acids degradation and folate biosynthesis.

### 3.4 Transcriptional landscape of *Tetradesmus obliquus slm1*

We compared changes in gene expression between the starchless mutant (*slm1*) and WT to understand the differences resulting from the lack of starch accumulation. An overview of gene expression data in *slm1* in comparison to WT is shown in figure 1C. The PCA shows very similar regulation over time between both strains. While many time points display similar expression patterns, there are also clear differences related to the separation in time phases. At the time points from 16h until 3h (phase IIIa to phase Ia), all *slm1* samples along PC1 are shifted left compared to WT, but they remain at the same level along PC2. This reflects a state of expression of WT that is never reached by slm1, rather than a diurnal dysregulation. On the contrary, the samples at 6h and 9h along PC2 are shifted up for *slm1* as compared to WT, but remain at the same level along PC1. This reflects an early state of expression, especially for *slm1* samples at 6h, which fits into the phase IIb. Another smaller time dysregulation is a small shift of time point 22h that is shifted down for slm1 as compared to WT. This reflects an early state of expression that fits into to the time phase Ia. The relative time phases of *slm1* are also shown in figure 1.E. Overall, two main trends are observed: earlier changes in expression of processes shortly after light and a delay in change of expression of the processes before the dark period.

The PCA considering the whole set of expressed genes (Supplementary file S2) displays a very similar expression between WT and *slm1*. While there is a general overlap, the expression of *slm1* seems noisier, especially for the time points between 19h and 0h.

#### 3.4.1 Comparison of the gene expression dynamics in WT and *slm1*

Using the regression based approach of maSigPro, we identified genes with differences in expression profile between *slm1* and WT. Our experimental design included sequencing samples of WT every hour and of *slm1* every three hours. However, maSigPro is designed to allow uneven distribution of time points. 784 genes showed no significant differences in expression profile between WT and *slm1* strains. 3902 genes showed significant differences in expression profile between WT and *slm1*. Additionally, 40 genes were found to have a time profile in the *slm1*, while no time profile was detected in WT for these genes. Apart form having no time profile it could also be that these genes are too noisy or are not expressed in WT.

#### 3.4.2 Differences in time profiles

Clustering was done for the gene expression data obtained from the mutant *slm1*. Analysis of clustering performance indicators again resulted in 6 clusters. As previously stated, 784 genes did not show any significant difference in time dynamics between the two strains, therefore, these genes were used to guide the identification of the mutant clusters in relation to the wild type clusters. The cluster assignments of these 784 genes in both strains is shown in figure 3A. The genes in WT clusters 3 and 4 are all found in the same *slm1* cluster, consequently named 3&4. Likewise, the WT genes in clusters 1 and 6 are all found in the same *slm1* cluster, consequently named 1&6. The WT genes in cluster 2 and 5 ended up in separate *slm1* clusters and consequently kept the same cluster name. Finally, the two remaining clusters, named A and B, do not contain genes with conserved expression. Those clusters have a time profile that does not fit any of the cluster profiles in WT.

**Figure 3:**
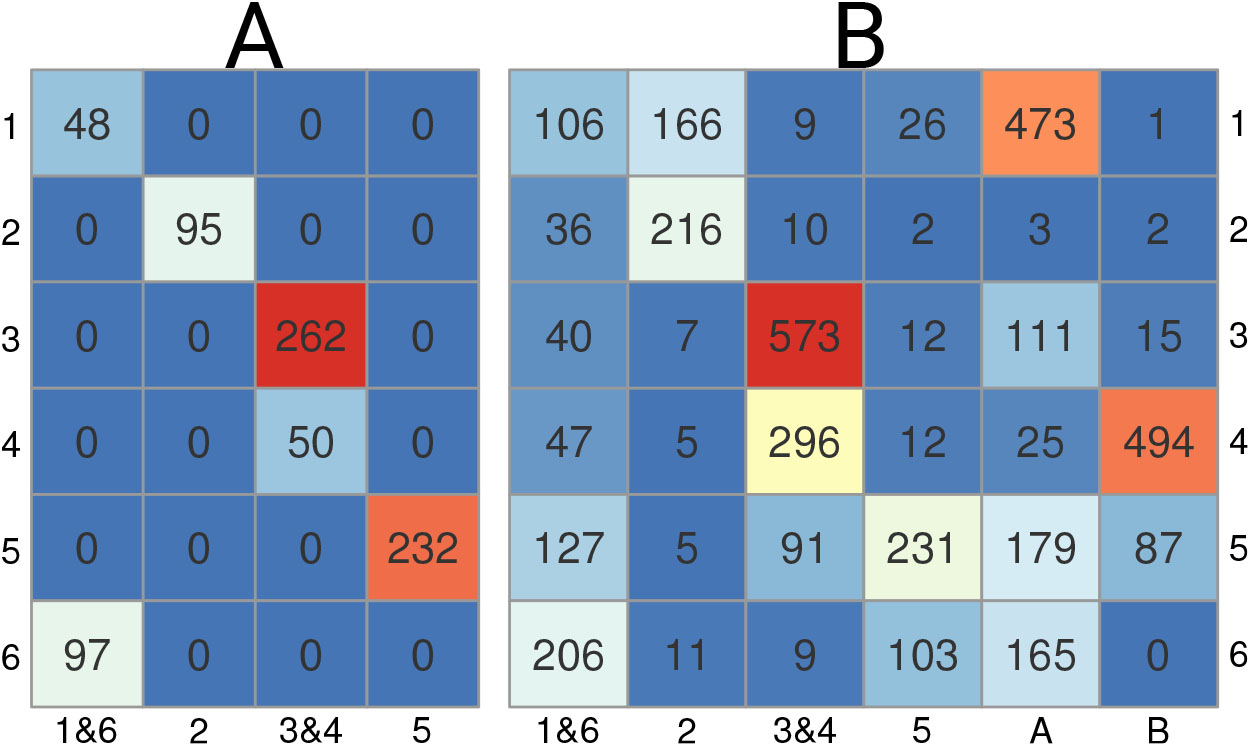
Distribution of genes between time profile clusters of Tetradesmus obliquus wild-type (WT) (rows) and slm1 (columns). A: 784 genes with the same time expression profile in slm1 and WT. B: Genes with significantly different time profiles between slm1 and WT, as identified by maSigPro (3902 genes). Background color ranges from blue, white, to red, with red being the highest value and blue the lowest.

Figure 3B shows the cluster assignments of the 3902 genes with significant differences in their time profiles when comparing WT and *slm1*. From these genes with altered expression (Figure 3B), there is a general trend of genes to remain clustered together. This means that most genes are transcriptionally controlled in large groups. The genes in WT cluster 3 are almost all found in the associated *slm1* gene cluster 3&4. Again, almost all genes in WT cluster 2 are found in the associated *slm1* cluster 2. The genes in WT cluster 5 are distributed over five *slm1* clusters, with the majority in the associated *slm1* cluster 5. The genes in WT cluster 6 are distributed over three clusters, with the majority in the associated *slm1* cluster 1&6. Interestingly, a minority of genes in WT cluster 4 is found in the associated *slm1* cluster 3&4, but the majority is found in *slm1* cluster B. Similarly, a minority of WT genes in cluster 1 is distributed over three *slm1* gene clusters, with a small portion in the associated *slm1* cluster 1&6, a bigger portion in *slm1* cluster 1, and the majority in *slm1* cluster A. Thus the new cluster A mainly receives genes from the WT clusters 1, and to lesser extent from cluster 3, 5 and 6. On the other side, the new cluster B receives genes mainly from cluster 4 and to a lesser extent from cluster 5.

*slm1* cluster median profiles are depicted in Figure 4, together with the associated WT clusters. From cluster 1&6 to cluster B each *slm1* cluster grouped respectively 735, 507, 1304, 621, 958, and 600 genes. Overall, the time profiles of *slm1* clusters show strong similarities with the associated WT clusters. *slm1* clusters A and B are displayed together with WT clusters 1 and 4, respectively, due to the large number of genes found in common. The new cluster A is characterized by low expression at dark and the first half of the light period, with a single expression peak during the second half of the light period. The new cluster B is characterized by a shift in the timing of the two peaks and a change in amplitude. A large number of genes were detected to be expressed differently between the two strains, but are associated to clusters with similar profile. These genes are more affected on their amplitude, and not necessarily on the time regulation.

**Figure 4:**
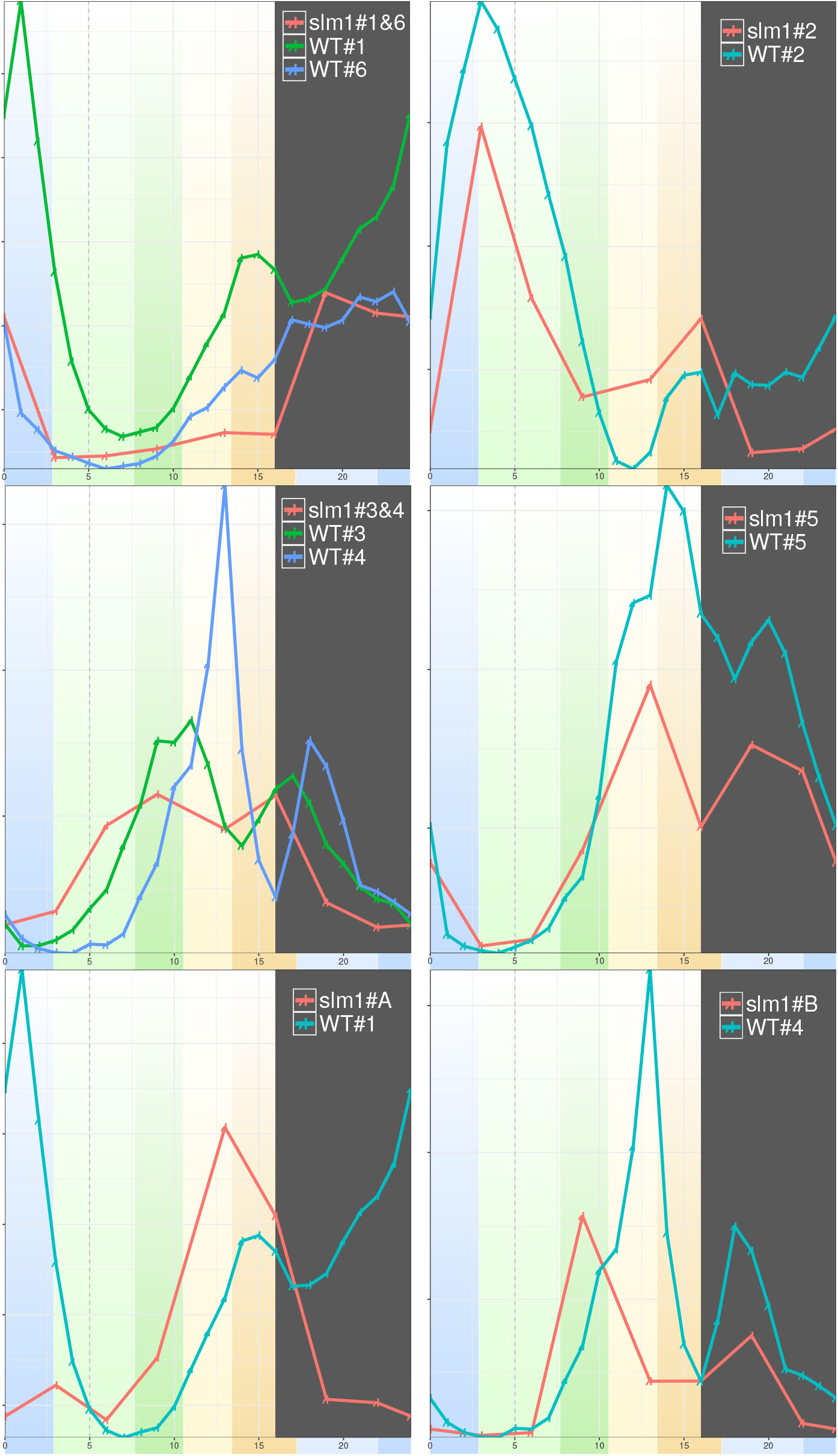
Expression profiles of clusters obtained with slm1 samples and their associated profiles in the WT. WT clusters 1 and 6, slm1 cluster 1&6; WT and slm1 cluster 2; WT clusters 3 and 4, slm1 cluster 3&4; WT and slm1 cluster 5; cluster A; WT cluster 4 and slm1 cluster B. Slm1 clusters are plotted with the related WT clusters according to the common conserved genes and the redistribution of genes as in Figure 3. The background colors correspond to the colors given to the time phases in Figure 1. The gray area represents the dark period.

91 genes that were found in WT cluster 5, are now found in *slm1* cluster 3&4. This represents a shift in time that is reflected in an earlier expression of these genes. This correlates with the observed early expression of genes along PC2 in phase II. A substantial number of genes from WT cluster 1 are found in *slm1* cluster 2, suggesting that these genes are shifted towards later expression in time and are not expressed during the dark period anymore. Again, the transfers of genes from WT clusters 4 and 5 to *slm1* cluster B reflect an earlier change of expression. Opposite to the previously mentioned changes, which all indicate decreased expression during dark, the expression of genes in *slm1* cluster 1&6 are now expressed exclusively during the dark period.

#### 3.4.3 Functional differences between gene clusters in *slm1* and WT

As observed by the similarities and changes of expression between the two strains (figure 3B), genes have related expression in large groups. As a result, the enrichments are consistent with the large groups of genes changing expression or not. The detailed enrichments are available in supplementary file S4. Enrichments of *slm1* clusters A and B are respectively similar to WT clusters 1 and 4. *slm1* cluster 3&4 is not enriched for the two AA pathways found in WT cluster 3 (lysine degradation and phenylalanine metabolism). This cluster also remains enriched in cellular structure and DNA replication, but without any biological process GO terms. Unlike for WT cluster 6, *slm1* cluster 1&6 do not display enrichment for other fatty-acids related pathways. AA synthesis and protein translation activities are now enriched in *slm1* cluster A, but were enriched in WT cluster 1. In *slm1*, nitrogen metabolism is no longer found enriched in cluster 5. This is due to three EC numbers (out of the five) that are associated to *slm1* cluster A. Globally, the enrichments are following the expectations drawn from the gene transfers described in Figure 3B.

## 4 Discussion

This study analyzed gene expression dynamics in a 16:8h light-dark (LD) cycle synchronized culture of *T. obliquus* wild-type (WT). Hourly sample collection and RNA sequencing allowed us to observe in detail the changes of gene expression during a diurnal cycle. A starchless mutant of the same species (*slm1*) was also studied. *Slm1* was cultivated in the same conditions, but RNA samples were taken every three hours. The succession of cellular events were then compared between WT and *slm1*, allowing us to examine the role and importance of starch as a transient energy storage compound in this microalga.

### 4.1 The diurnal rhythm of *T. obliquus wild-type* is driven by six transcriptional phases

#### 4.1.1 Transcriptional phases

In this experimental setup, *T. obliquus* cells are synchronized to the diurnal light-dark (LD) cycles [8]. Many organisms synchronized their metabolism to anticipate the changes in the environment [9, 45, 46]. For photosynthetic microorganisms, this synchronization provides a benefit as they can capture sunlight efficiently during the day and perform light sensitive processes at night [6, 12]. In T. *obliquus*, synchronization was observed in growth and cell division, as well as in changes in biomass composition [8]. The overall analysis of the changes in gene expression in both WT and *slm1* strains shows a circular pattern in the PCA plot over two strong principal components. In WT, changes in expression appeared to occur sequentially, back and forth along each PC. In the studied process, two main effects impacting gene expression are light availability and time itself.

By correlating the samples (figure 1B and 1D) and correlating genes expressions (clusters), we identified six time phases and six clusters of genes, respectively. The agreement between these two independent analyses indicates that they are related to each other. In fact, the time phase changes correspond to jumps in expression between the time points, when either large increase or large decrease in overall gene expression occurs. The gene clusters correspond to synchronized peaks of expressions. Thus, the identified phases are associated to peaks in expression for each clusters. Accordingly, cluster 1 peaks during phase Ia; cluster 2 during IIa; cluster 3 during IIb, cluster 4 during IIIa, cluster 5 during IIIb, and, finally, cluster 6 during Ib. Because the cluster regroup genes from the same biological processes, the six transcriptional phases correspond to a logical succession of biological processes.

#### Temporal succession of cellular events as described by the gene clusters

An overview of the changes in the transcriptional landscape and experimental measurements of WT through the diurnal LD cycle is presented in Figure 5.

**Figure 5:**
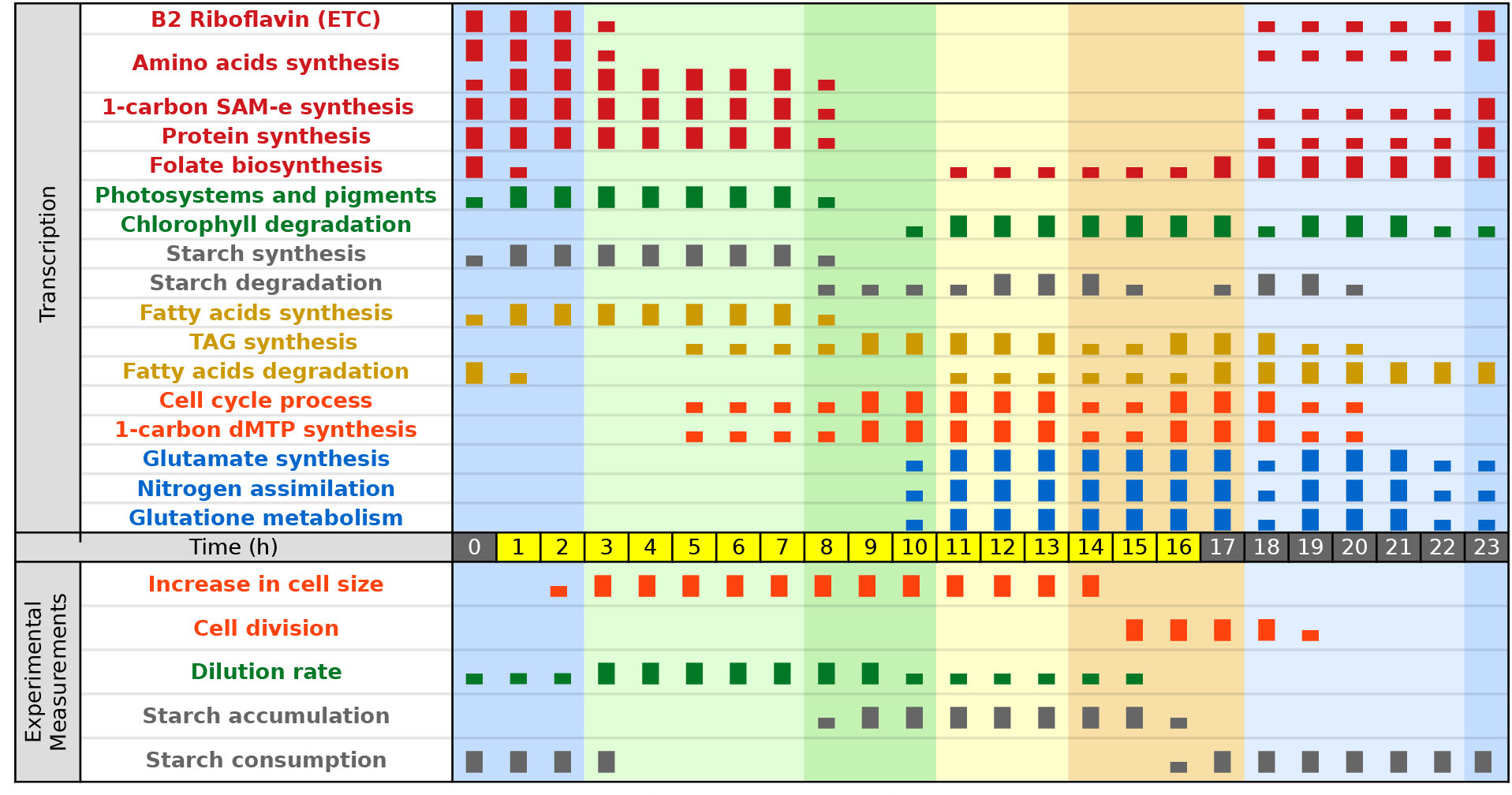
Overview of the diurnal changes in the transcriptional landscape and experimental measurements of T. obliquus wild-type under 16:8h light/dark cycles. Experimental measurements were taken from [8]. Small and big rectangles indicate that the process is occurring at relatively low or high level, respectively. Groups of related processes are indicated in the same color. The background colors correspond to the colors given to the time phases in Figure 1. Amino-acids biosynthesis two lines reflect the synthesis of different amino-acid fitting different cluster profiles.

The median profile of the genes in cluster 1 peaks the first after light. The enrichment of cluster 1 reflects amino acids (AA) and protein synthesis. At the same time, riboflavin metabolism is found enriched and its product FAD is an important reducing cofactor for cellular respiration. Additionally, the pathway ubiquinone and other terpenoid-quinone biosynthesis in enriched in cluster 1. The overview of the pathway (supplementary file S5) indicates that ubiquinone is the most probable product in this time frame.

Ubiquinone is an important step in the oxidative phosphorilation by oxidating FADH2 back into FAD. As such, it indirectly plays a role in ATP synthesis. Overall, these are strong signs of catabolism of either starch or fatty acids for the production of energy (ATP). This is in agreement with the observed use of starch at the beginning of the day. The small peak of expression found around 15h is probably a result of energy requirements for cellular division and night processes. Other AA synthesis pathways are also found enriched in different clusters, but cluster 1 contains the majority.

Cluster 2 mainly contains genes involved in pigment and starch synthesis, besides carbon fixation and AA synthesis. The main contributing genes in the PCA (Figure 1A) were also found in cluster 2. These main contributors are the light harvesting complexes, which are known to display large variations in transcription. The expression profile of this cluster matches the dilution rate profile, where the gene expression precedes the dilution rate with about 3 hours [8].as can be seen in Figure 5. This time delay between gene expression and formation of the actual light absorbing complexes is as expected. In cluster 2, among the proteins associated to carbon fixation, GAPDH was found to be a main contributor in PC2. Three of the four reactions involved in the starch synthesis were found in cluster 2, this can be observed from the pathway map (supplementary file S5). Detailed analysis is described below in a dedicated section. The expression profile of processes related to starch synthesis and degradation also show good correlation to the starch measurements (Figure 5), again, taking into account a certain delay between expression and appearance of the phenotype. Starch is a transient energy source and is subject to simultaneous synthesis and degradation [8, 47], which results in daily fluctuations between net production during the day and consumption during the night and early day. Expression patterns of the starch synthesis genes match the observed starch accumulation. Between 16h and 0h, only degradation occurs presumably to provide energy for the night processes, since light is not available. Between 0-4h, starch is still consumed. During this phase, starch is apparently still used to supply additional energy and carbon for the build up of new photo systems, which is also up regulated at this time. Between 4-8h, starch synthesis increases following the up regulation of the starch synthesis genes in cluster 2. The starch synthesis machinery is getting fully operational and energy demanding processes, such as AA and protein synthesis, are down regulated, resulting in net starch synthesis.

Following cluster 2, the genes in cluster 3 show a peak in expression at 13h. This cluster is enriched for genes involved in the cell cycle and protein maturation. There seems to be an important level of protein maturation through glycosylation in the endoplasmic reticulum at this stage. Furthermore the GO enrichment shows terms associated with progression through the cell cycle, meaning that DNA replication probably starts in the following hours. Finally, GO terms on organelle fission, chromosome organization, and organelle organization, point towards cell division which agrees with the observed cell division that starts just before the night and continues for a main part during the early night.

Shortly after the genes in cluster 3 peak, genes associated to cluster 4 increase in expression. This cluster contains genes related to degradation of various glycans together with enrichment in several carbohydrate metabolic pathways. The information on individual genes indicates that these reactions are taking part in starch degradation. Starch is degraded during the night, which is thus in agreement with the expression of these genes just before the start of the night in cluster 4. Furthermore, the large diversity of enrichment in carbohydrate related pathways is suggesting inter-conversion between carbohydrates.

The next cluster for which the gene expression shows a peak is cluster 5. Cluster 5 also shows enrichment for carbohydrates pathways, but more importantly it is enriched in glycolysis. ATP synthase is found in this cluster too, but the gene is highly expressed throughout the whole light period with highest expression before dark. Together, they clearly indicate that the genes for starch degradation and glycolysis are providing the means for the organism to generate the needed energy during the night. Glutamate associated genes are also found in cluster 5. Glutamate synthesis is analyzed in a dedicated section for nitrogen assimilation. Glutamate synthesis, which is at the center of nitrogen assimilation and amino-acid metabolism, seems to be upregulated before and during the night as anticipation for the synthesis of AA (cluster 1) and pigment (chlorophyll and carotenoids, cluster 2) during the night and early light period. It also suggests that *T. obliquus* continues to assimilate nitrogen during the night, which is in agreement with the observed nitrate consumption during the night by WT (data not shown). Notably, *slm1* does not consume nitrate in the night and nitrogen metabolism is not enriched for *slm1*, which suggests that pathways in nitrogen metabolism are regulated by the energy status of the cell. However, nitrate reductase is not annotated and nitrite reductase has extremely low expression with no time regulation. Cluster 5 also shows an overall increase of expression in genes associated to glutathione reductive cycle, indicating some form of oxidative stress. This stress could be a side effect of the whole day exposure to light leading to accumulation of reactive oxygen species and damaged photosystems. Among the genes associated to the chlorophyllase (EC 3.1.1.14), the best candidate (g19660.t1) is found in cluster 5. This also agrees with the enrichment of Cytochrome P450, which was recently found to take major part in the breakdown of chlorophyll in *Arabidopsis thaliana* [48–50].

Finally, cluster 6 is enriched in fatty acid degradation, which appears to occur during the night and early morning. However, fatty acid degradation was not observed experimentally. Possibly, these enzymes are involved in remodeling of membranes that has to occur after cell division, or maybe are involved in recycling unsaturated fatty-acids damaged from oxidative stress.

Folate (vitamin B9) is a vital cofactor, notably in thymidylate (dTMP) synthesis and in S-adenosylmethionine (SAM-e) synthesis. The gene expression in the pathway “one carbon pool by folate” of WT (supplementary file S5) is clearly decoupled for these two processes. While dTMP related reactions are associated to cluster 3, SAM-e reactions are associated to clusters 1 and 2. This result is also displayed in Figure 5, in which their respective synchronization with cellular division and protein synthesis is clear.

Interestingly, “folate biosynthesis” was found enriched in cluster 6 (Table 2), together with the reactions from GTP until the precursor of dihydrofolate (DHF). Considering the temporal succession of these events, it seems that folate is being accumulated before the light period in order to sustain the high levels of protein synthesis at dawn.

In summary, the annotation and enrichment analyses uncovered the general function of the genes in each cluster, thus showing a clear succession of cellular events. Overall, the gene clusters represent the following general functions, by order of highest peak appearance: RNA transcription, gene expression, protein synthesis, pigments synthesis, starch synthesis, fatty acids synthesis, protein glycosylation, cellular growth and division, starch degradation and sugars inter-conversions, riboflavin synthesis, folate synthesis, and, finally, fatty acids degradation (Figure 5). This succession of transcriptional events and their relative expression in time indicates that *T. obliquus* does not simply adapt its cellular phenotype to direct signals from the light switch, but it also anticipates the changes in light conditions. For example, before the light is on, the microalga starts synthesizing important metabolites that will fulfill the high demand of gene expression and protein synthesis. This allows quick synthesis of the photosystems and storage of energy and carbons into starch. Likewise, starch degrading enzymes are upregulated before the night starts to ensure the supply of energy and carbons required for building up AA during the night. This indicates some kind of time keeping mechanism.

Green algae are known since 1950s for performing cell division over multiple fission. *C. reinhardtii* and *Scenedesmus quadricauda* were studied, and models of their multiple fission cell cycle were established [51, 52]. *T. obliquus* is from the same family as *S. quadricauda*, hence, their cell cycle should be very similar. By comparing our enrichment analyses and the established cell cycle model, it is logical to think that the processes leading to the cell growth (G1), DNA replication (pS), mitosis (G2), and cellular division (G3), are driven by genes found in cluster 2, 3, 4, and 5, respectively. Relating to our time phases, they are correlating to phases IIa, IIb, IIIa and IIIb, respectively. In our experimental condition, the doubling time was measured to be 0.67 day in WT and 0.75 day in *slm1*. Meaning that WT divides 1.5 times a day, and *slm1* divides 1.33 times a day. This can be explained by a double fission for a portion of the cells, 50% for WT and 33% for *slm1*. Looking at the time profiles, we found cluster 3, 4 and 5 to have a double peak of expression 4 to 5 hours apart, with significantly lower expression at the second peak. This relatively lower expression at the second peak could be associated to that portion of cells performing a double fission.

### 4.2 Impact of starch deficiency on the diurnal rhythm

When analyzing data from *slm1*, significant changes in the temporal profiles were identified for the vast majority of the genes compared to the WT (3902 genes). In most cases, the genes remained clustered together, as for the WT, and most differences can be explained by a time shift or a change in amplitude. Time changes of expression in phase II contains most of the overall differences between WT and *slm1*. It is possible that phase IIIa is also affected by some difference in time regulation, but three out of four samples of *slm1* 13h are aligned along PC1 with the other night samples. This is an example where the less frequent sampling of *slm1* (every 3h) may conceal important changes in expression and have impacted the detection of expression changes and the cluster separation. From the six clusters of WT, two clusters (3-4 and 1-6) could not be efficiently separated in *slm1*. However, two extra clusters of time profiles were found: A and B. Whereas cluster B could be confounded with WT cluster 4, cluster A showed a really novel expression profile not seen in WT. The profile in cluster A in *slm1* shows a very low expression during the night and the beginning of the day, with a single peak in expression occurring during the second half of the day. On the other hand, in WT, the profile of these genes more or less matches the observed net consumption of starch. In WT net starch consumption occurs during the night and early day, which may be associated to the expression of the genes appearing in cluster A in the mutant *slm1*. Or in other words, impossibility to consume starch during this period in *slm1* may be related to the low expression of the genes occurring in cluster A.

The biochemical data for *slm1* also match the transcriptome results, but to a lesser extent than for WT. Analysis of data obtained from the starchless mutant *slm1* shows that the phases from the WT are to some extent preserved. However, some time shifts are observed.

Starch deficiency seems to result in rescheduling biological processes from dark to light period. It is possible that these processes could not happen at night due to the lack of energy originally provided by starch degradation. Processes required for expression and translation were transferred from cluster 1 to cluster A in *slm1*. In cluster 1, the expression rises before and during the dark period to peek at the beginning of the day, whereas in cluster A, expression is high before dark and stays low during the dark period. Thus, expression of these genes seems to depend on the presence of starch and are possibly related to the energy status of the cell. In *slm1*, the expression is shifted to the second half of the light-period, which is normally the time when starch accumulates and for which it could be assumed that sufficient energy is available. Glutathione metabolism was found enriched in cluster A, but it does not reflect a strong change in expression compared to WT in cluster 5, besides the absence of expression during the dark period.

Inspection of the profile of *slm1* cluster 1&6, shows that genes in this cluster are mainly expressed during the dark period. This indicates that the scheduling of dark period processes not depending on energy sources only suffers small changes. Notably, genes associated to faty acid degradation in *slm1*, are in this cluster. This points to TAG and lipid degradation occurring during the dark period, although no significant reduction in the TAG content was observed.

### 4.3 Selected reactions and pathways

In addition to the described time shifts caused by starch deficiency, a number of processes show differences in expression between *sml1* and WT that seem more complex. These are discussed in the following paragraphs.

#### 4.3.1 Starch synthesis

The diurnal expression for the genes related to the reactions analyzed in this section are depicted in Figure 6. While enrichment for “starch and sucrose metabolism” was associated to cluster 3, the detailed view at the pathway (supplementary file S5) revealed that the reactions causing the enrichment are related to starch degradation and sugar interconversions. More interestingly, three of the four enzymes necessary for starch synthesis are found enriched in cluster 2, being the ADP-Glucose pyrophosphorylase (EC:2.7.7.27), the starch synthase (EC:2.4.1.21) and the granule-bound starch synthase (EC:2.4.1.242). The fourth, starch branching enzyme, was not found to be regulated in time. Gene expression associated to these reactions are displayed in Figure 6. Two genes were associated to the starch synthase (g7327.t3 and g2865.t1). Their expression profiles in WT are complementary, each being highly expressed during one half of the light period. However, their expression is altered in *slm1*, with expressions that appear stretched during the light period. This could be the response to the lack of ADP glucose. The granule-bound starch synthase was associated to different clusters for each strain, but their profiles are very similar. However, just like the starch synthase, the granule-bound starch synthase expression in *slm1* appears stretched over the light period. We also noticed that the expression of this enzyme peaks in the middle of the light period. The difference in time regulation of the two type of starch synthase could be explained by their associated processes on starch branching nature [53]. The amylose isomerase (EC:2.4.1.18) was associated to two genes, with very similar profiles, but without apparent time regulation for either strain.

**Figure 6:**
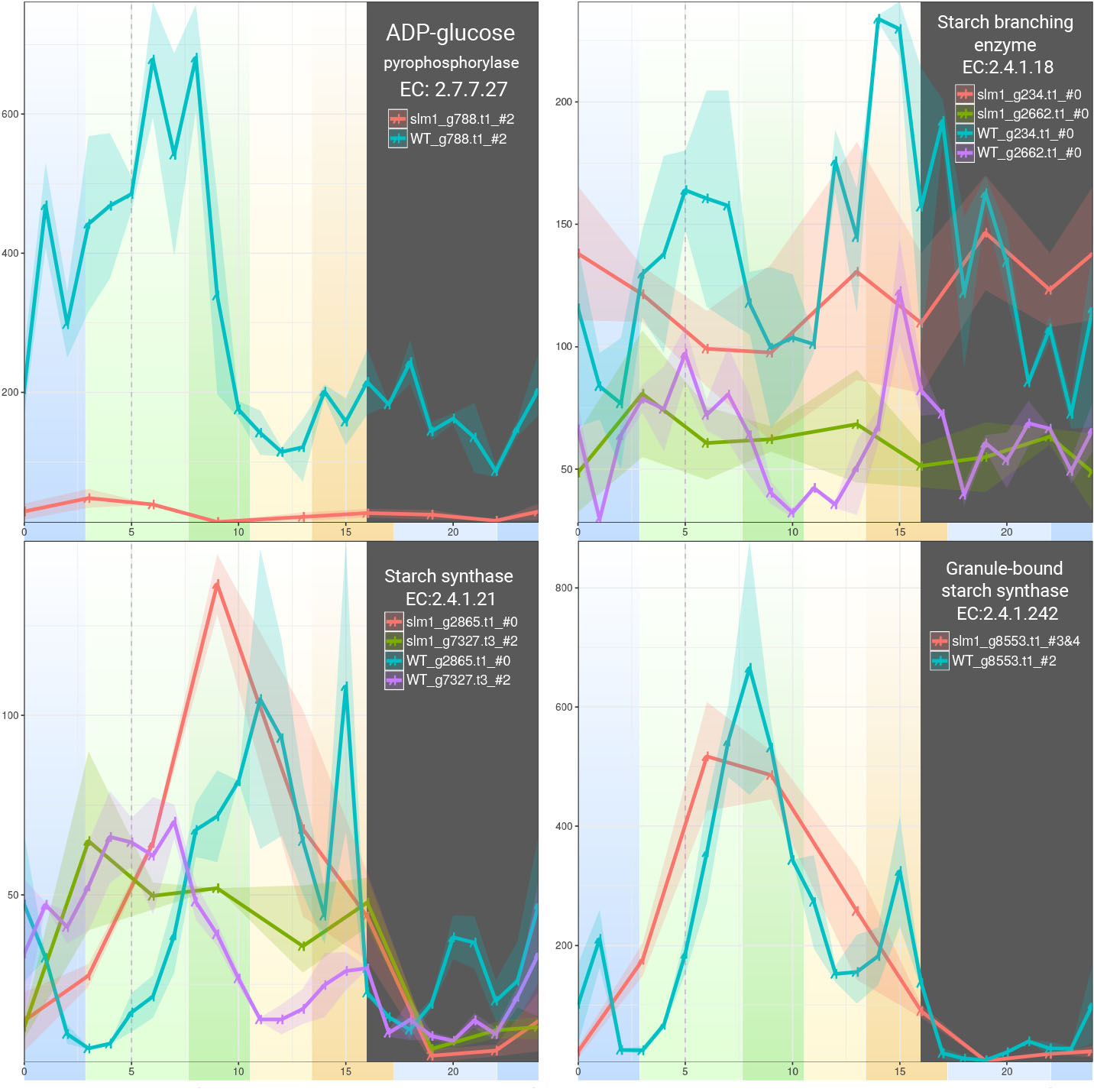
Diurnal expression of genes related to carbon fixation. The points represent the mean for each time points, the ribbon covers the minimum and maximum values for each time points. The genes associated to the same EC number are plotted together. All genes in the same plot are using the same y axis scale (not shown), but the scale between each of the six plots varies. Each plot is labeled with the reaction name, the corresponding EC number, and the color legend for each gene-strain combinations. For each gene-strain combination, the letter or the number behind the # symbol, indicate the cluster with which the gene is associated. The background color corresponds to the time phases from figure 1. The dark gray area corresponds to the dark period.

The reason of the incapacity of *slm1* to synthesize starch has already been studied [17]. The gene coding for the small subunit of ADP-glucose pyrophosphorylase was found to contain a non-sense mutation. In conditions of continuous light, this gene was found to be strongly down-regulated in comparison to WT (approx. 5 folds). In this study, we found that this gene (g788.t1) is even more strongly down-regulated, meaning that the changes become even more prominent in presence of a diurnal cycle. Additionally, we found that the time regulation is preserved between the two strains. On the other hand, the big subunit of the ADP-glucose pyrophosphorylase was annotated to two genes with identical protein sequence (g14915.t1, g9029.t1), but their expression is much lower than the small subunit (approximately 500 folds). Although their expression was too noisy to be selected by maSigPro, their expression pattern in WT is very similar to the small subunit. (high expression during the first 10h of the light period). In *slm1*, their expression during the light period is noisier and seems lower than WT.

#### 4.3.2 Carbon fixation

The diurnal expression of the genes related to the reactions analyzed in this section are depicted in Figure 7. The maSigPro analysis indicated no significant temporal regulation of RuBisCO (EC:4.1.1.39) and phosphoglycerate kinase (EC:2.7.2.3). The expression of RuBisCO (Figure 7) is noisy, which explains the undetected time profile. Its WT expression seems a little higher around 0h and around 9h, and lower right before dark. In s*lm1*, expression of RuBisCO seems generally lower during the light period. The logical explanation is that less carbon can be fixed due to starch deficiency. For the bisphosphatase, the WT profile is noisy but with a clear pattern of higher expression between 4h and 9h. In *slm1*, the bisphosphatase expression does not display the higher expression observed in WT. Other reactions associated to carbon fixation are fitting the expression pattern of clusters 1 or 2 in both WT and *slm1*. Among those reactions, the two enzymes preceding in the carbon fixation pathway (phosphoriboisomerase and phosphopentokinase) are co-localized with the RuBisCO in the pyrenoid [54]. These enzymes follow an expected pattern with carbon fixation during the light with a strong peak at the beginning of the light period.

**Figure 7:**
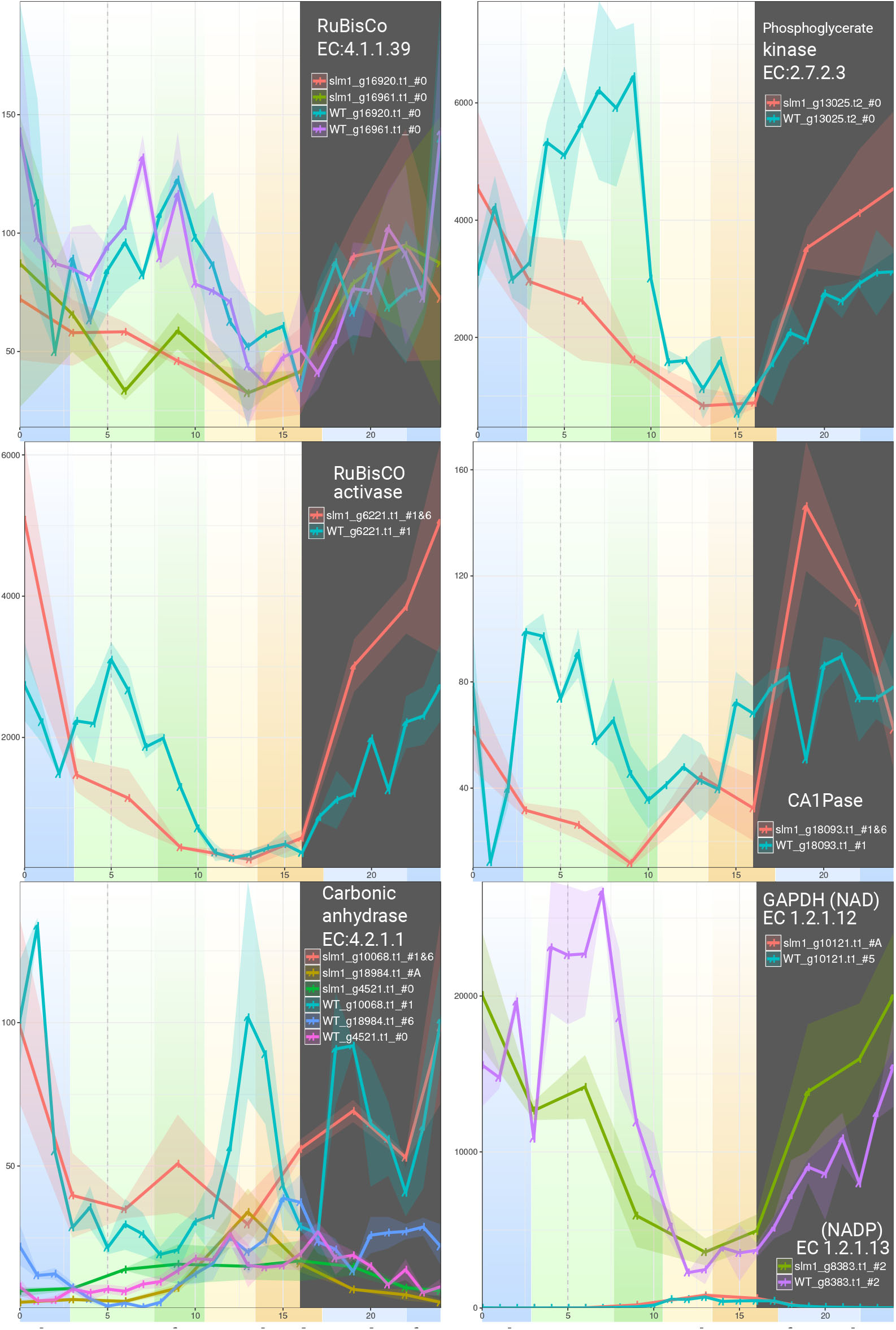
Diurnal expression of genes related to carbon fixation. The points represent the mean for each time points, the ribbon covers the minimum and maximum values for each time points. The genes associated to the same EC number are plotted together. All genes in the same plot are using the same y axis scale (not shown), but the scale between each of the six plots vary. Each plot is labeled with the reaction name, the corresponding EC number, and the color legend for each strain_gene_cluster combination. For each gene-strain combination, the letter or the number behind the # symbol, indicates the cluster in which the gene was found. The background color corresponds to the time phases from Figure 1. The dark gray area corresponds to the dark period.

Due to the low affinity of RuBisCO to CO2, algae maintain high concentrations of CO2 around the RuBisCO. This is known as the carbon-concentrating mechanisms (CCM). Carbonic anhydrases (CA, EC:4.2.1.1) are used to convert CO2 into bicarbonate to minimize diffusion through the membranes. A specific CA (Cah3) was found to be co-localized with the RuBisCO to perform this task [55]. It was also found that this enzyme helps to maximize PSII activity via the redox potential provided by the bicarbonate. Our analysis detected two genes following a time pattern that were associated to CA function. DELTA-BLAST reveals one with a high sequence similarity to the aforementioned Cah3 protein from *Chlamydomonas reinhardtii* (g18984.t1), and the second is closest to the cytosolic CAs Cah8 and Cah7 of the same species (g10068.t1). As can be seen in Figure 7, the *T. obliquus* candidate gene for Cah3 (g18984.t1) is displaying a time profile associated to WT cluster 6, with higher expression during the end of the day and night. This enzyme needs to be active from early hours after light for the PSII to benefit from it. Interestingly, its expression in *slm1* is following the profile of cluster A, meaning expression exclusively before dark. The other CA gene is associated to Cah8 of *C. reinhardtii* (g10068.t1) and is found in *slm1* cluster 1 due to its peak of expression at 1h, but with additional peaks at 13h and during the dark period. Its expression is relatively higher than Cah3, and its regulation is similar between the two strains.

The RuBisCO activase is activated by light and helps to release the substrate from the RuBisCO active site, accelerating the reaction [56, 57]. No EC number is associated to this protein, but we found three candidate genes, among which only g6221.t1 displays high expression and a detected time profile. It is found in WT cluster 1 and displays some increasing expression during the night with a peak at 0h. Interestingly, its expression remains high during the first half of the light period. In *slm1*, it shows a peak of expression during the night, that is higher than in WT and then it gradually drops but remains high during the first hours of the day. Overall, the expression of the RuBisCO activase resembles the expression of the RuBisCO, but with higher amplitude and with few hours earlier.

In complement to the RuBisCO activase, the RuBisCO inhibitor 2-Carboxy-D-arabitinol 1-phosphate (CA1P) is inactivated by the 2-carboxy-D-arabinitol-1-phosphatase (CA1Pase EC: 3.1.3.63). CA1P normally binds to RuBisCO under dark conditions, which prevents it from performing chemical reactions. We found that in WT, the gene g18093.t1, associated to the CA1Pase, displays a rather constant expression through both light and dark periods, except for a dip in expression around 1-2 h. However, in *slm1*, CA1Pase has a higher expression during the dark period and lower during the day.

All the aforementioned enzymes are co-localized in the pyrenoid with the RuBisCO. This sub-cellular micro-compartment plays a major role in carbon fixation and it is logically localized in the chloroplast, along the thylakoid membrane. It is found in green algae (chlorophyta), and therefore found in *T. obliquus* [58].

Pyrenoids are not delimited by a membrane, but they accumulate starch at their periphery in the form of sheath. It is believed that starch is a barrier that limits the transfer of CO2 to the rest of the chloroplast and outer compartments. It was demonstrated that the pyrenoid plays a role in the CCM, allowing higher levels of carbon fixation in *C. reinhardtii* [59], however analysis of a *C. reinhardtii* starchless mutant showed that the starch sheath surrounding the pyrenoid is not involved in the CCM [60]. Therefore, the starchless mutant, *slm1*, should not suffer from a lessened CCM in comparison to the wild-type. The lower yield on light should be due to the lack of a transient energy storage allowing for harvesting more light energy during the day and using this in the night.

#### 4.3.3 Nitrogen metabolism

The diurnal expression of the genes related to the reactions analyzed in this section are depicted in the Figure 8. Nitrogen metabolism was found enriched in WT cluster 5, but this enrichment is lost in *slm1*. Inspection of KEGG metabolic maps (supplementary file S5) shows that the enrichment in WT is due to five EC numbers found in cluster 5: 1.7.99.1, 1.4.7.1, 1.4.1.14,1.4.1.3, and 1.4.1.4. For *slm1*, the genes associated to the first three of these EC numbers are found in cluster A.

**Figure 8:**
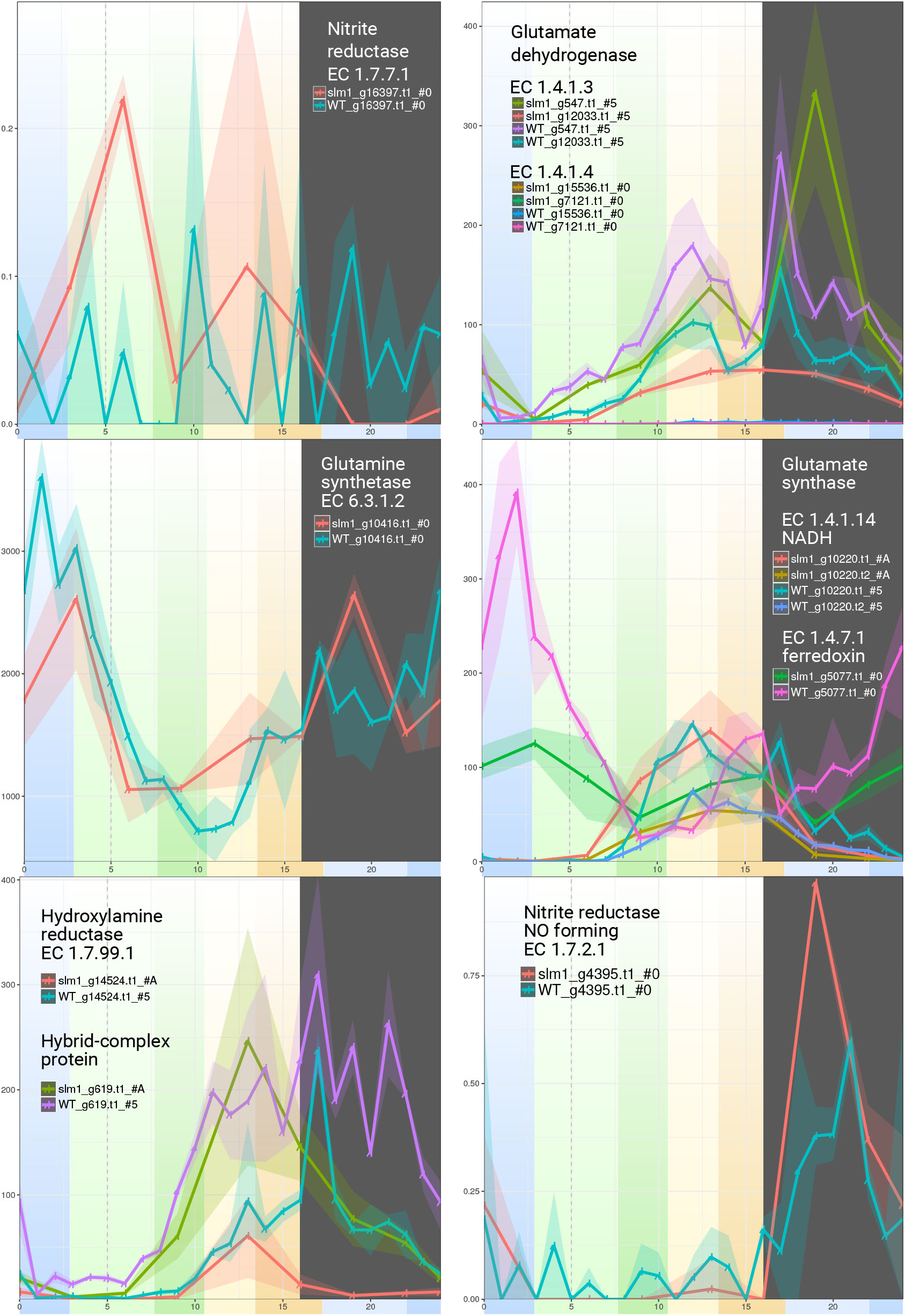
Diurnal expression of genes related to nitrogen assimilation. The points represent the mean for each time points, the ribbon covers the minimum and maximum values for each time points. The genes associated to the same EC number are plotted together. All genes in the same plot are using the same y axis scale (not shown), but the scale between each of the five plots vary. Each plots are labeled with the reaction name, the corresponding EC number, and the color legend for each strain_gene_cluster combinations. For each gene-strain combination, the letter or the number behind the # symbol, indicate the cluster in which the gene was found. The background color corresponds to the time phases from Figure 1. The gray area corresponds to the dark period

The first enzymes of the nitrogen assimilation pathways are nitrate and nitrite reductases. They are known to be strongly regulated by the light and nitrogen condition of the cell [61]. Nitrate reductase in green algae is located in the pyrenoid [54]. We were not able to identify the corresponding gene in the transcriptome. We identified the nitrite reductase gene and it displayed extremely low expression (with zero expression for few time points) and no identifiable diurnal profile. Nitrate and nitrite reductase convert nitrate to ammonium. Ammonium can be transformed into glutamate via two pathways, either by the glutamate dehydrogenase (GDH, EC 1.4.1.3 and 1.4.1.4), or via the glutamate synthase cycle that consists of glutamine synthase (GS, EC 6.3.1.2) followed by glutamate synthase (EC 1.4.1.14 and 1.4.7.1). GS is known as the main contributor for nitrogen assimilation [62], whereas GDH is generally involved in balancing nitrogen between metabolites by recycling ammonium for GS.

The candidate gene for GS (g10416.t1), which is known to be located in the chloroplast [62], was also associated to the chloroplast by PredAlgo [63]. This matches with the observed, yet noisy, higher expression at night and early day. In *slm1*, the expression is very similar and at the same level of expression to WT, still, the expression during the dark period seems to peak at the same time as the GDH (g547.t1). For the glutamate synthase, we have three candidates, two using NAD as a cofactor, and one using ferredoxin as a cofactor. The two transcripts associated to the NAD-dependent reaction originate from the same gene and their expression profile is very similar with low expression at the beginning of the day and peaking at the end of the day and decreasing again during the night. The WT expression of the candidate associated to the ferredoxin-dependent reaction is very similar to GS but with a small time-shift (approximately 2h) towards later expression. Additionally, we also notice a small peak of expression right before the dark period and comes right after the two genes associated to the NAD-dependent glutamate synthase. Additionally, the ferredoxin-dependent glutamine synthase also peaks right after the GDH during the late light period, but also several hours after the GDH peak during the dark period. These observations in WT match the observed behavior of these enzymes in *C. reinhardtii* [64], meaning that the ferredoxin-dependent enzyme, cannot only perform the assimilation during the light period, but also contributes to the assimilation during the dark period. Expression of the ferredoxin-dependent enzymes in *slm1* follows a similar temporal profile but is strongly reduced.

A gene was identified for each of the GDH g547.t1 and g12033.t1 associated to EC 1.4.1.3 and 1.4.1.4 respectively. Both are in cluster 5 and have similar WT expression profiles. In *slm1*, g547.t1 exhibits a similar profile with the second peak shifted to two hours later. Expression of g12033.t1 in *slm1* appears to have lost sharp peaks with very gradual variation. The opposing expression of GDH and GS matches previous observations in *C. reinhardtii* [64], suggesting GDH to be an exclusively catabolic enzyme. These profiles suggest nitrogen recycling at the end of the light period and beginning of dark. Differences in the GDH are probably related to differences in cofactor utilization [61]. Nitrogen concentration measurements (data not shown) revealed that *slm1* does not assimilate nitrogen during the dark period, which would suggest EC 1.4.1.4 as the main route for nitrogen fixation during the dark period.

Nitrite reductase (NO forming, 1.7.2.1) is expressed almost exclusively during the dark period in both strains with higher expression in *slm1* (see Figure 8). Our annotation associated two genes to the hydroxylamine reductase. Among these, g14524.t1 is associated to EC 1.7.99.1 and evidence from *Chlamydomonas* suggest that hydroxylamine might function as a late intermediate in the catalytic cycle of nitrite reductase [65]. The other one, g619, appears to be a hybrid-cluster protein (HCP, best matching HCP3 of *C. reinhardtii*). HCP is considered unique due to its iron-sulfur-oxygen complex and is known to have a very diverse range of functionalities such as hydroxylamine reductase activity and nitrite oxide reductase [66, 67]. In WT, hydroxylamine reductase is highly expressed starting from half of the light period (9h) and during the dark period. Similarly, HCP is progressively expressed at the end of the light period, but peaks immediately after dark and decreases progressively until the light period. In *slm1*, both hydroxylamine reductase and HCP are affected by starch deficiency, leading to reduced expression at dark. This is the strongest effect from the mutation in the nitrogen metabolism, it also correlates with the lack of nitrogen consumption during the night in *slm1* [8].

## 5 Conclusions

Oleaginous microalgae are a promising source of biofuels. Among the oleaginous microalgae, *Tetradesmus obliquus* is a promising candidate that can reach a high maximum TAG yield on light and TAG content under nitrogen starvation. However, in order to develop strategies to enhance lipid productivity for commercial production, a system-level understanding of metabolism is essential. Large scale microalgal production will be done outdoors under natural LD cycles. Therefore, the impact of LD cycles has to be carefully considered when characterizing the behavior of microalgae.

In this work, we show that LD cycles induce systems level transcriptional changes in 4686 genes showing a clear expression pattern and indicating a strict succession of cellular events. These cellular events were found to be in accordance with the biochemical measurements. While some regulations seem a direct response to light, other regulations reflect an anticipation to the switch from light to dark and vice-versa. This observed anticipation indicates an inner time-keeping system.

Additionally, we studied the diurnal transcriptional changes in the starchless mutant of *T. obliquus slm1*. Pthways directly associated to energy storage, such as carbon fixation, and other processes such as nitrogen metabolism are strongly affected in *slm1*. This cleavage of processes is attributed to starch deficiency, which seems to be the only transient energy storage compound in *T. obliquus*. Genes associated to TAG and lipid degradation are highly expressed during the dark period in *slm1*. This suggests TAG as a transient energy storage, however, no significant changes in TAG content were measured. As a result of missing source of energy during the dark period, *slm1* is less prepared to start the new cycle right after dark. More subtly, processes related to the cellular cycle were detected to start earlier in *slm1* than in WT.

Overall, we provide for the first time a diurnal transcriptional landscape under LD cycles with a high resolution (1h intervals) of an oleaginous green microalgae that produces both starch and TAG under nitrogen starvation. This is also the first time a diurnal transcriptional landscape is described and compared for a starchless mutant of green algae. The diverse set of insights revealed in this analysis of the diurnal cycle of *T. obliquus* is very valuable to develop strategies to increase yields. We suggest that the presented transcriptional landscape should be carefully considered when designing future experiments with LD cycles, including metabolic engineering approaches.

## Funding

This research project was supported by the Consejo Nacional de Ciencia y Tecnología – CONACYT, Mexico, Scholar 218586/Scholarship 314173. In addition, GMLS is part of the program “Doctores Jóvenes para el Desarrollo Estratégico Institucional” by the Universidad Autónoma de Sinaloa. This work has been financially supported by the Systems Biology Investment Program of Wageningen University, (KB-17-003.02-29).

## Acknowledgments

The authors would like to thank Tom Schonewille for the RNA extraction of all samples.

## Competing interests

The authors declare that they have no competing interests.

## Supplementary Files

**S1: Annotation.** The association between gene, protein, EC number, GO terms is summarized in this table sheet. The protein sequences are also given in this file.

**S2: PCA and heatmap of all expressed genes, before time profile selection.** In the PCA of WT, the two first component explain a big majority of the variations, and the time points follow a clear ordered circular pattern. The PCA with both strains displays a more noisy pattern and the

**S3: Finding optimal hierarchical cluster separation.** Two methods were used to find the optimal separation of genes into clusters. The first figure contains a plot for each index calculated for every cluster separation steps. The second figure contains the heatmap representation of the Rand index values generated by comparing cluster separation one on one. All these indexes were generated from R package “cluster.stats”.

**S4: Cluster enrichment.** Results of the enrichment analyses for each clusters of each strains, in both pathways and GO terms. Separate enrichment are also displayed for the gene with conserved expression between strains and those with altered expression, described in table 1 and figure 3.

**S5: Images of cluster colored KEGG pathway maps.** Compressed archive file of every annotated maps in PNG format. EC numbers colored according to related genes cluster association. Each EC number box is divided in seven parts: the first for genes with no time regulation and therefore no cluster association, the following six to the clusters defined in the strain. The colors and positions are depicted in the legend file called “_0_color_legend.png”. For WT, the colors are the same as given in figure 2. For *slm1*, the colors are given based on their WT counterpart when possible. Gene not regulated in time, filtered out by our analysis and by extend not clustered were added on the maps as gray color.

